# Toxicity and applications of internalised magnetite nanoparticles within live *Paramecium caudatum* cells

**DOI:** 10.1101/109488

**Authors:** Richard Mayne, James Whiting, Chris Melhuish, Andrew Adamatzky

## Abstract

The nanotechnology revolution has allowed us to speculate on the possibilities of hybridising nanoscale materials with live substrates, yet significant doubt still remains pertaining to the effects of nanomaterials on biological matter. In this investigation we cultivate the ciliated protistic pond-dwelling microorganism *Paramecium caudatum* in the presence of excessive quantities of magnetite nanoparticles in order to assess both potential beneficial applications for this technique as well as any deleterious effects on the organisms’ health. Our findings indicate that these nanoparticles are well-tolerated by paramecia, who were observed to consume in quantities exceeding 10% of their body volume: cultivation in the presence of magnetite nanoparticles does not alter *P. caudatum* cell volume, swim speed, growth rate or peak colony density and cultures may persist in nanoparticle-contaminated medium for many weeks. We demonstrate that *P. caudatum* cells ingest starch coated magnetite nanoparticles which facilitates their being magnetically immobilised whilst maintaining apparently normal ciliary dynamics, thus demonstrating that nanoparticle biohybridisation is a viable alternative to conventional forms of ciliate quieting. Ingested magnetite nanoparticle deposits appear to aggregate, suggesting that (a) the process of being internalised concentrates and therefore detoxifies nanomaterial suspensions in aquatic environments and (b) *P. caudatum* is a candidate organism for programmable nanomaterial manipulation and delivery.

## 1 Introduction

There are two major justifications for research into the hybridisation of nanoscale materials with biological matter. Firstly, despite their widespread use in recent years, significant doubt remains as to the potential deleterious effects of nanoparticles and nanomaterials on biological matter as even inert materials may be rendered reactive, immunogenic or otherwise harmful to life when fabricated in nano-scale quantities [1]. Secondly, nanomaterials may be fabricated to exhibit highly desirable characteristics (electrical properties, magnetism, high tensile strength etc.), the conferral of which to live cells would potentially lead to next-generation technologies such as bio-computer interfaces for restorative and/or augmentative medical applications.

Magnetite (iron II/III oxide) nanoparticles (MNPs) (a variety of super-paramagnetic iron oxide nanoparticles, SPIONs) are one such nanomaterial possessing desirable properties for applications involving live cells. Exhibiting superparamagnetism and apparently low toxicity, SPIONs are now routinely used as a contrast medium in *in vivo* magnetic resonance imaging and are a proposed drug-delivery (or otherwise therapeutic) agent for cancer treatment [2, 3]. With regards to their effects on other protists, the same class of nanoparticle may be hybridised with the plasmodium of slime mould *Physarum polycephalum* towards enhancing its value in bio-computer interfaces and unconventional computing devices [4–7]. Recent evidence has suggested that MNPs are not as biocompatible as once thought, however, due to their potential for bioaccumulation and generation of reactive oxygen species within host cells [8, 9]. Furthermore, little is known of the ecotoxicological significance of SPIONs released into the environment but recent studies on water-dwelling eukaryotes such as *Daphnia* spp. and plant life have demonstrated that they may potentially disrupt aquatic ecosystems [10].

This study examines the ciliated protistic pond organism *Paramecium caudatum* being cultivated in the presence of excessive quantities of MNPs in order assess the potential uses of nanohybridised paramecia whilst concurrently observing for any deleterious effects on the organisms’ health. We conclude by discussing the apparent effects of this treatment on *P. caudatum* cells and the ecotoxicological significance of these results, and present several novel applications for hybridised *P. caudatum* cells containing nano-magnetite.

## 2 Materials and Methods

*P. caudatum* cultures were cultivated in Chalkley’s medium enriched with 10 g of alfalfa and 40 wheat grains per litre. Cultures were exposed to a day/night cycle but were kept out of direct sunlight; ambient temperatures ranged from 19–24 *°*C.

In experiments where *P. caudatum* cells were exposed to MNPs, suspensions of 200 nm (hydrodynamic diameter) starch matrix-coated multi-core MNPs (Chemicell GmBH, Germany) were added to fresh culture medium at a concentration of 0.25 mg ml^−1^ (approximately 2.2×10^12^ particles per ml). This concentration was chosen as a comparable quantity of nanoparticles per unit biomass to our previous studies with other single-celled organisms [5]. Stock cultures at a concentration of approximately 1000 cells ml^−1^ were harvested in log growth phase and added to the nanoparticle-infused culture medium, in which they were incubated for periods in excess of two months.

The following microscopical measurements were made on a regular (daily) basis (n = 3 per culture per day for each): average cell count, average swimming speed, total cell cross sectional area and percentage of cross sectional cell area occupied by cytoplasmic/vesicular inclusions whose colour was suggestive of MNP deposits. Cells were observed in glass microscope well slides. Cells were chemically fixed in order to record photomicrographs. Fixation was achieved by adding 10 *µ*l of 4% paraformaldehyde in pH 7.0 phosphate buffered saline solution to each slide well after swim speed measurements had been made.

Observations were made with a Zeiss Axiovert 200M inverted microscope and photo/videomicrographs were captured with an Olympus SC50 digital camera via CellSens software, in which brightness and contrast adjustments were made.

For cell volume measurements, each image was analysed to extract both the cells’ cross sectional area in squared micrometers, but also to estimate the percentage of internalised MNPs. Each image was imported into Matlab (Mathworks, USA) and processed in the following manner: the image was converted into greyscale, and a threshold was applied to extract all material darker than the background media. Each threshold was visually checked in order to ascertain that only organisms were isolated in each image. The number of pixels isolated by the threshold was then summed to give the cells’ total cross sectional area. This process was also performed using a different threshold to determine the number of pixels whose colour value corresponded to dark cytoplasmic inclusions in order to identify any internalised MNPs. As the threshold values had to be determined visually for each image separately, this process was performed ‘blind’, i.e. without the operator knowing whether each image were a control or test measurement. The number of pixels isolated by this method were then compared with the cells’ total volume to give a percentage value.

Video analysis for measurement of organism swimming speed was also performed using Matlab. RGB images were imported from the video frame-by-frame for sequential analysis and organism positioning. For each video set, the organisms were isolated from the RGB image by colour, whereupon the data for each frame was converted to a JPEG image file for further analysis. To detect the position of the organisms, a Laplace template of a Gaussian filter was defined before being convolved over the image; the size of the filter was iteratively determined by visual feedback of the user. After organism detection on every frame had occurred, the positional data was passed to a bespoke Kalman filter which accurately estimates the position of the particle across each frame using the data from the full time-series of particle positions to predict and confirm the movement of each organism. From this it is possible to measure the speed of each organism in a noisy video, creating a dataset of organism speed and momentary position. While the script ran, frame-by-frame images were shown on screen allowing visual validation of positional tracking by the authors. Average speed was calculated for each organism.

All numerical data were subject to statistical analysis in Matlab: two-tailed t-tests and Mann-Whitney U tests were used.

## 3 Results & Discussion

All measurement data are shown in Table 1. The microscopical appearance (morphology and swimming patterns) of *P. caudatum* cells treated with MNPs was not noticeably altered aside from the inclusion of rust-coloured deposits and more dark objects within intracellular vesicles and the cells’ cytoplasm (Fig. 1). Organisms treated with MNPs contained approximately 5% more dark intracellular inclusions than controls. No statistical difference was observed in total cell volume or swimming speed between controls and test organisms.

**Table 1.** Table to show mean (*x̅*) and standard deviation (*σ*) values for measurements of total cell cross-sectional area (*xa*, in *µm*^2^), cell content comprising dark objects (*doc*, in percentage), swim speed (*ss*, in *µm*^2^), growth rate (*µ*, in h^−1^) and peak colony density (*pcd*, in cells ml^−1^). Asterisks indicate a statistically significant difference in means to controls at p *<* 0.0001 (T-test and Mann-Whitney U-test)

| | $xa \bar{x}$ | $xa \sigma$ | $doc \bar{x}$ | $doc \sigma$ | $ss \bar{x}$ | $ss \sigma$ | $\mu \bar{x}$ | $\mu \sigma$ | $pcd \bar{x}$ | $pcd \sigma$ |
| --- | --- | --- | --- | --- | --- | --- | --- | --- | --- | --- |
| Control | 12715.66 | 5994.59 | 7.33 | 1.67 | 132.54 | 42.79 | 0.020 | 0.009 | 309 | 120 |
| Test | 13381.09 | 4700.77 | 12.68* | 3.16 | 144.94 | 31.64 | 0.021 | 0.009 | 317 | 123 |

**Fig. 1.**
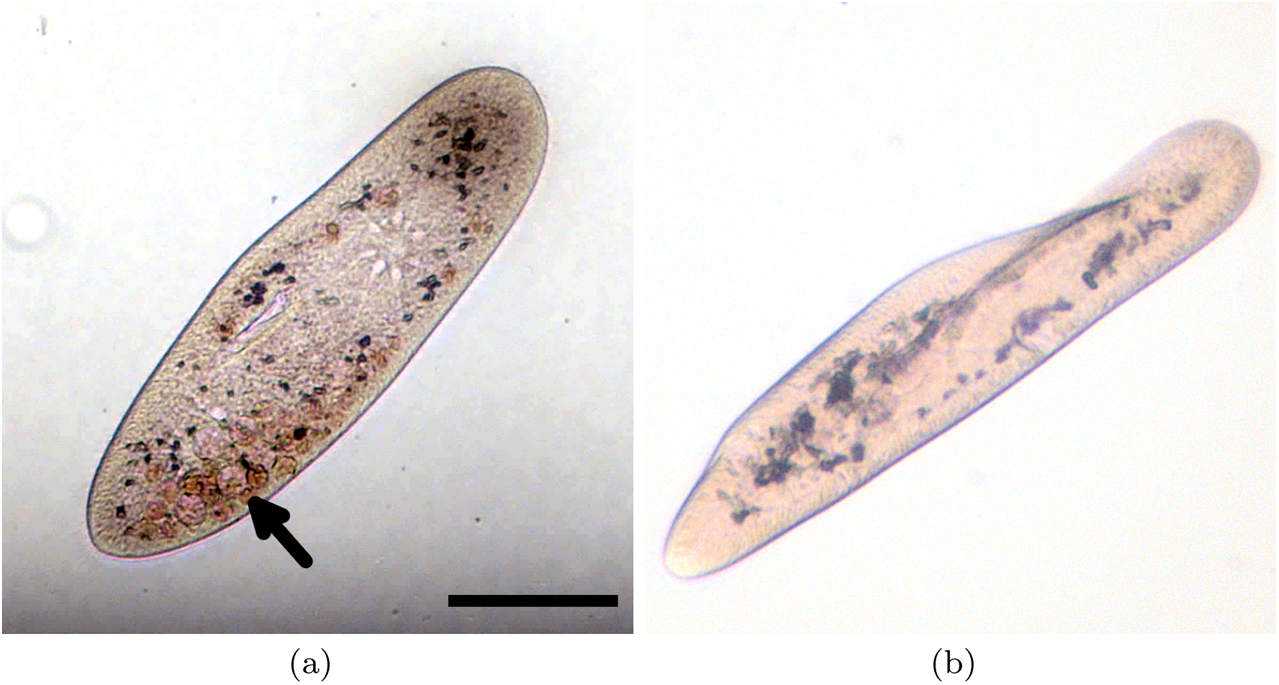
Photomicrographs to show appearance of *P. caudatum* cells (unfixed): (a) Following exposure to MNPs. Multiple rust-coloured deposits (arrowed) are visible in the cytoplasm. (b) Control, demonstrating a lack of rust-coloured deposits. Scale bar 100 *µ*m.

Furthermore, no statistical difference in growth rates or peak colony density were observed between controls and organisms treated with MNPs. All cultures (test and control) persisted over the duration of the experiment (14 days).

Our results indicate that the *P. caudatum* cell is highly tolerant to being cultured in the presence of large quantities of MNPs, as indicated by our not observing any deleterious effects on the health of the organisms with regards to their size, morphology, motility, growth rate or colony density. This indicates that, despite recent evidence suggesting that this variety of nanoparticle may be harmful to aquatic microorganisms, they do not appear to induce any readily-observable toxicological effects in our ciliated model organism.

Intriguingly, the observations of MNPs within the organisms suggests that they were ineternalised in the manner of a nutrient source, noting that the size of the individual particle cores, 10 nm, were too large to enter the cell via any other route such as through membrane pores. This apparent uptake of MNPs was likely a result of their starch coating. Interactions between *P. caudatum* and MNPs with other coatings/no coating remain a topic for further study.

Although no measurements of MNP intracellular reactivity (e.g. generation of reactive oxygen species) or the longevity of individual cells were made, the longevity of cultures treated with MNPs was not significantly different from that of controls as both varieties were kept in culture for periods exceeding two months (data not shown).

We propose that *P. caudatum* may detoxify certain environmentally-dispersed nanomaterials: that dark/rust-coloured deposits could be easily identified in *P. caudatum* cells indicates that they are aggregated *in vivo* into non-nanoscale objects. This reduction in surface area to volume ratio likely renders the deposits less reactive and therefore less harmful.

This apparent lack of toxicological effects incident of internalising quantities of MNPs allows us to speculate on the potential applications of this process of biological–artificial hybridisation. In further experiments, we exposed 10 *µ*l droplets on glass microscope well slides containing approximately 5 *P. caudatum* cells treated with MNPs to a 1.28 T 25x40x4.0 mm neodymium magnet. Holding the magnet in close proximity to the margins of the well caused the organisms to be drawn to the edge of the droplet where they were held immobile (Fig. 2). By increasing the distance between the magnet and margins of the well, the organisms were able to move but at a significantly reduced speed.

**Fig. 2.**
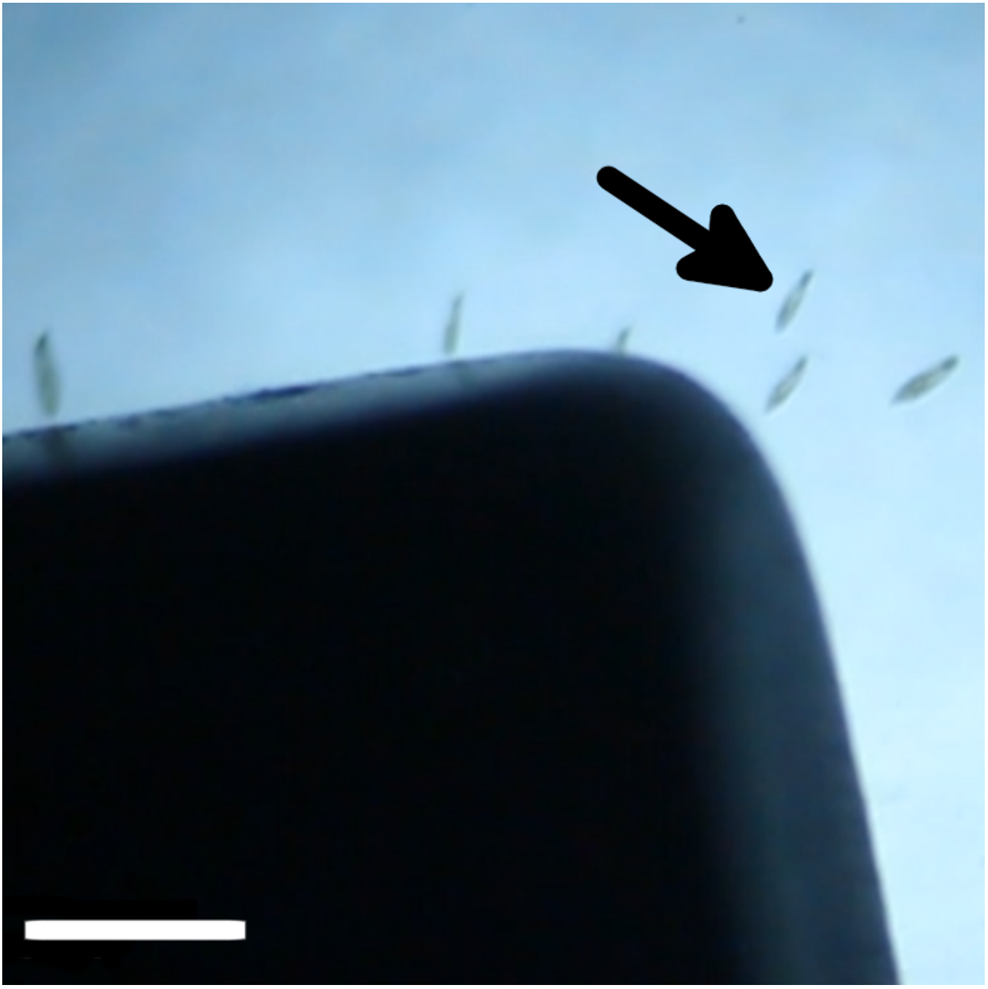
Stereomicrograph showing MNP-treated *P. caudatum* cells (arrowed) being attracted towards a permanent magnet (black object). Scale bar 500 *µ*m.

## 4 Conclusions

Magnetic restraint of ciliates via their hybridisation with biocompatible magnetic nanomaterials would appear to be an attractive alternative the established methods of microorganism immobilisation/quieting, such as: replacement of media with inert viscous fluids, induction of hypoxia through hermetically sealing the observation environment [11, 12], addition of low-concentration toxins (e.g. aliphatic alcohols [13], anaesthetic compounds [14]), ultraviolet light irradiation [15], establishment of fluid pressure gradients/microfluidic compression [16] and adhesion to solid surfaces via positively-charged proteins (3-aminopropyltriethoxysilane, protamine sulphate) [17].

The advantage of magnetic restraint over these other quieting methods is that it does not necessitate inducing deleterious health effects on the organism, maintains the chemical composition and hence physical characteristics of the fluid medium (thus minimising interference with natural ciliary beating processes) and allows for momentary adjustment of the strength of attraction (i.e. by moving the magnet or using magnets of different strengths). Although magnetic restraint of *Paramecium spp.* has been previously described via internalised magnetite (particles of ca. 3 *µ*m diameter), the authors did not describe the use of microparticles in this context as being a method for fully immobilising the organisms [18].

Finally, this ciliated model organism’s capacity for gathering, internalising and concentrating nanomaterials holds exciting possibilities for the prospect of orchestrated biological manipulation and delivery (guided by gradients of attractants, repellents or magnetic fields) of nano and micro-scale compounds of interest, although further research is required in this area before practical applications can be realised.

